# *In vivo* multiphoton fluorescence imaging with polymer dots

**DOI:** 10.1101/227488

**Authors:** Ahmed M. Hassan, Xu Wu, Jeremy W. Jarrett, Shihan Xu, David R. Miller, Jiangbo Yu, Evan P. Perillo, Yen-Liang Liu, Daniel T. Chiu, Hsin-Chih Yeh, Andrew K. Dunn

## Abstract

Deep *in vivo* imaging of vasculature requires small, bright, and photostable fluorophores suitable for multiphoton microscopy (MPM). Although semiconducting polymer dots (pdots) are an emerging class of highly fluorescent contrast agents with favorable advantages for the next generation of *in vivo* imaging, their use for deep multiphoton imaging has never before been demonstrated. Here we characterize the multiphoton properties of three pdot variants (CNPPV, PFBT, and PFPV) and demonstrate deep imaging of cortical microvasculature in C57 mice. Specifically, we measure the two-versus three-photon power dependence of these pdots and observe a clear three-photon excitation signature at wavelengths longer than 1300 nm, and a transition from two-photon to three-photon excitation within a 1060 – 1300 nm excitation range. Furthermore, we show that pdots enable *in vivo* two-photon imaging of cerebrovascular architecture in mice up to 850 μm beneath the pial surface using 800 nm excitation. In contrast with traditional multiphoton probes, we also demonstrate that the broad multiphoton absorption spectrum of pdots permits imaging at longer wavelengths (λ_ex_ = 1,060 and 1225 nm). These wavelengths approach an ideal biological imaging wavelength near 1,300 nm and confer compatibility with a high-power ytterbium-fiber laser and a high pulse energy optical parametric amplifier, resulting in substantial improvements in signal-to-background ratio (>3.5-fold) and greater cortical imaging depths of 900 μm and 1300 μm. Ultimately, pdots are a versatile tool for MPM due to their extraordinary brightness and broad absorption, which will undoubtedly unlock the ability to interrogate deep structures *in vivo*.

## Introduction

*In vivo* imaging with multiphoton fluorescence microscopy (MPM) is widely used due to its ability to provide diffraction-limited, three-dimensional images of intact tissue at depths ranging from a few hundred microns to more than 1 mm.^1–7^ Recent reports of deep tissue, *in vivo* imaging with two- and three-photon excitation have demonstrated the potential of imaging structures in the brain beyond the cortex, including the corpus callosum and hippocampus.^8,9^ These advances have been driven in part by the availability of new ultrafast laser sources at longer wavelengths (λ_ex_ = 1000 – 1600 nm) than those traditionally used in two-photon fluorescence imaging (λ_ex_ = 700 – 900 nm). Equally important to the availability of laser sources is the availability of contrast agents with strong multiphoton absorption properties at these longer wavelengths. In this paper, we show that an emerging class of fluorophores termed polymer dots^10^ possess strong two- and three-photon excitation across a broad range of wavelengths and enable deep imaging of microvasculature in the brain.

MPM of vasculature requires the intravenous injection of bright, biocompatible contrast agents that preferably exhibit large absorption cross sections under near-infrared (NIR) excitation (λ_ex_ = 700 – 1000 nm) and prolonged blood circulation times. Traditional exogenous contrast agents for *in vivo* MPM include organic dyes such as dextran-conjugated fluorescein and indocyanine green or inorganic semiconductor quantum dots.^11,12^ Organic dyes, however, suffer from poor photostability and low quantum yields in aqueous biological environments.^13.14,15^ Although quantum dots offer improved brightness and photostability, they present substantial toxicity concerns and are prone to bioaccumulation in organs and tissues.^16,17,18^ Thus, the biological imaging community is eager for a safer, brighter, and more stable probe for deep, high-resolution *in vivo* imaging.

The highly fluorescent semiconducting polymer dot (pdot) is a promising candidate for *in vivo* MPM with material properties that can potentially overcome many of the limitations faced by other probes^19^. Although pdots are similar to quantum dots with respect to size (5–50 nm) and quantum yield, pdots are brighter, more photostable, and present no clear evidence of biotoxicity.^19,20^ A useful measure of fluorescence brightness is the action cross section, which is given by the product of the peak absorption cross section and the fluorescence quantum yield.^19,21^ Polymer dots rival both quantum dots and organic dyes in that their two-photon action (2PA) cross-sections are one to two orders of magnitude larger than inorganic quantum dots and three to five orders of magnitude greater than commonly used fluorescent dyes.^19,22^ Herein, for the first time, we present evidence that we can take advantage of polymer dots’ many favorable properties to produce high-quality *in vivo* multiphoton images of vasculature with excellent signal-to-background ratio (SBR) at depths exceeding 1 mm.

Another advantage of pdots is their broad absorption, which enables multiphoton imaging with a variety of ultrafast laser sources, including ytterbium-fiber lasers (yb-fiber, λ_ex_ =1060 nm) and longer wavelength optical parametric amplifiers (OPA, λ_ex_ = 1100 – 1400 nm). Such broadband compatibility takes advantage of the favorable photophysical characteristics of these unique laser sources to improve SBR beyond the capabilities of conventional 2P titanium-sapphire (Ti:S) microscopy. Moreover, the ability to excite polymer dots at longer wavelengths allows us to approach an ideal biological imaging wavelength situated at 1,300 nm where absorption and tissue scattering events are minimized.^8,23^ For instance, the photophysical advantages of longer wavelength excitation of PFPV (λ_ex_ = 1,060 nm) coupled with the ytterbium-fiber laser’s intrinsic pulse characteristics results in a 3.5-fold improvement in SBR, and an overall 50 μm gain in penetration depth *in vivo*. Multiphoton imaging of CNPPV-labeled vasculature using an OPA (λ_ex_ = 1225 nm) increases SBR by ~8.2 fold (*z* = 700 μm), and extends imaging depth 450 μm further into the brain. Notably, PFPV and CNPPV both exhibit a partial three-photon power dependence at these longer wavelengths, which contributes to a larger SBR by the suppression of out-of-focus fluorescence.^24,25^ Overall, semiconducting polymer dots are ideal for MPM due their enhanced brightness over traditional fluorophores, and spectrally wide absorption range. These advantages coupled with their intrinsically low cytotoxicity help overcome the longstanding limitations of quantum dots and establish the immense potential of polymer dots for *in vivo* biological imaging.

## Results

### Two- and Three-Photon Excitation Power Dependence of Polymer Dots

Two-photon excitation of polymer dots has been successfully demonstrated by other research groups, although it has been more generally referred to under the larger and less specific umbrella of multiphoton imaging.^22^ With mounting evidence of the advantages of three-photon microscopy over 2P imaging for deep *in vivo* imaging, such as the suppression of out-of-focus fluorescence, reduced scattering, and improved SBR, the need to identify specific excitation wavelengths that produce two-versus three-photon absorption becomes imperative.^24–26^ Thus, we tested the excitation power dependence of various polymer dots including a poly[2-methoxy-5-(2-ethylhexyloxy)-1,4-(1-cyanovinylene-1,4-phenylene)] polymer (CNPPV) and two fluorene-based copolymers (PFBT and PFPV) at wavelengths ranging from 790 – 850 nm, 1,060 nm, and 1200 – 1350 nm (**Fig. S1**). We find that all three semiconducting polymers exhibit a strong two-photon dependence at 800 nm (**Fig. 1**). In the case of CNPPV and PFBT, this effect persists out to 1,060 nm whereas PFPV begins to demonstrate a partial three-photon power dependence at this wavelength (**Fig. S2**). Pure three-photon fluorescence is most strongly demonstrated at 1,300 nm for CNPPV, 1,350 nm for PFBT, and 1,325 nm for PFPV (**Fig. 1**). Within the 1,060 – 1,300 nm excitation range, there is a clear rising transition from two-photon to three-photon excitation. This data is especially instructive in that it serves as a reference for the selection of appropriate excitation sources and wavelengths for MPM experiments with polymer dots.

**Fig. 1.**
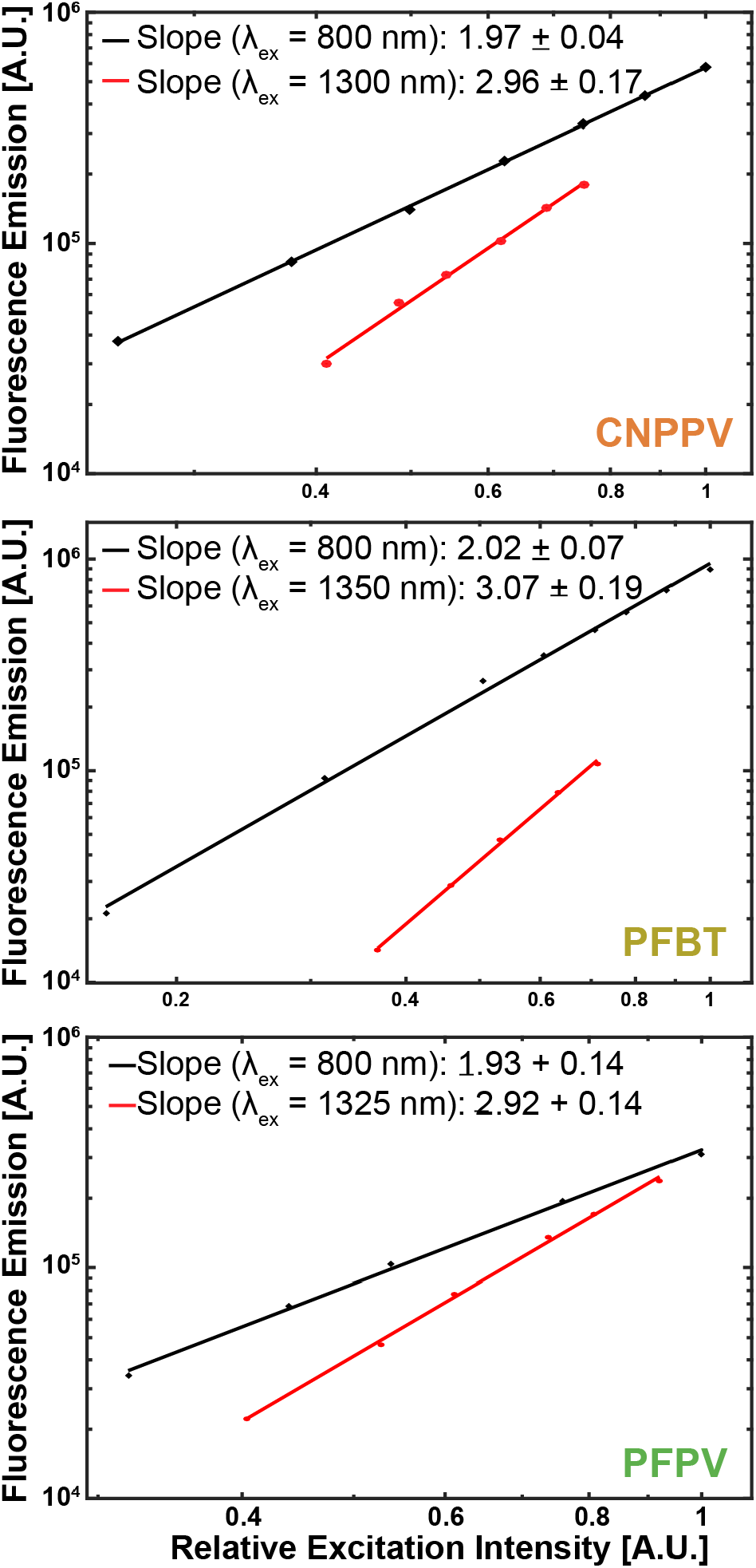
Logarithmic plots of the dependence of two- and three-photon induced fluorescence on excitation power. The excitation wavelength and fitted slope is indicated in the legend of each graph. The estimated uncertainty of each slope is reported as a standard error. Each plot corresponds to a distinct polymer dot species (*Top*: CNPPV, *Middle*: PFBT, *Bottom*: PFPV).

### Two-Photon Action Cross Sections of Polymer Dots

Characterization of polymer dot brightness at longer excitation wavelengths is a critical first step towards their adoption as probes for multiphoton microscopy. In particular, two-photon action (2PA) cross sections, given by the product of the peak two-photon absorption cross section (*σ*_2_) and quantum yield (*η*), provide a direct and exceptional measure of brightness. A comparison of MEH-PPV, a closely related variant of CNPPV, and PFPV’s 2PA cross-sections^22^ to quantum dots,^12,27^ organic dyes,^12,27^ and fluorescent proteins^27^ demonstrates that MEH-PPV and PFPV are the brightest among all fluorophores tested within a 770-870 nm excitation range with peak cross-sections of ~5.50×10^4^ GM and ~2.02×10^5^ GM, respectively (**Fig. S3**).^22^ In contrast, the reported 2PA cross sections of quantum dots range from ~2×10^3^ (QD630) to ~4.7×10^4^ GM (QD605) at these wavelengths.^12,28^ Organic dyes such as rhodamine B^27^ and fluorescein^27^ generally exhibit 2PA cross sections up to 5×10^2^ GM with typical values around 10 GM,^12^ and common fluorescent proteins such as YFP (8.63 GM) are often more dim.^27^ Thus, polymer dots are one to two orders of magnitude brighter than quantum dots and roughly three to five orders of magnitude brighter than conventional organic dyes and fluorescent proteins across the Ti:S spectrum.

### Two-Photon SBR is Improved by the Brightness of Polymer Dots

The primary goal of this paper is to evaluate the use of pdots for deep *in vivo* vascular imaging. C57 mice with intravenous injections of fluorescent dye, quantum dots, or polymer dots were imaged through an optical cranial window^29^ using 800 nm excitation. Dextran-conjugated fluorescein (λ_em_ = 524 nm; FD2000S, Sigma-Aldrich) served as the organic dye of interest, chosen because of its prevalent usage as a contrast agent in biological experiments.^21,28,30–32^ QD605 (λ_em_ = 605 nm; Q10001MP, ThermoFisher Scientific) was the selected semiconductor quantum dot, a probe known for its large quantum yield, atypically high brightness, and photostability. ^28,33^ Lastly, three polymer dot variants, PFBT (λ_em_ = 538 nm), PFPV (λ_em_ = 520 nm), and CNPPV (λ_em_ = 590 nm), were selected as they represent different classes of polymers, including poly(phenylene vinylene) and fluorene-based copolymers.^19^ Comparing 100 μm thick maximum intensity projections at the same cortical depths, the polymer dot data visually appears much brighter than the fluorescein and quantum dot images, demonstrating an enhanced SBR given by polymer dots (**Fig. 2**). A quantitative comparison reveals that PFPV produces the largest SBR, followed by PFBT then CNPPV (**Fig. S4**). Relative to QD605 and fluorescein, pdot SBR is larger throughout the entire depth range. To ensure fair comparisons, the average laser power at each depth was maintained at consistent levels across the separate imaging experiments. Furthermore, appropriate filters were selected for all experiments in order to maximize the collection efficiency of each fluorophore and ensure fair comparisons of the separate image stacks. Vasculature in mice labeled by QD605 (**Fig. 2**, *Right*) appeared noticeably distorted and discontinuous, and imaging was limited to a 550 μm depth relative to the cortical surface. This is explained by the fact that the mouse did not survive the retro-orbital injection of inorganic quantum dots due to toxic effects, resulting in an absence of active circulation to evenly distribute the contrast agent in plasma. Meanwhile, chronic *in vivo* imaging experiments with repeated intravenous injections of polymer dots in the same mice over the course of several months showed no signs of cytotoxicity or deleterious effects on the animals, supporting several published claims of polymer dot biocompatibility.^20,34–36^ An example of two-photon data collected from a mouse injected with 50 nM of PFPV showcases excellent SBR up to 650 μm deep and appreciable signal levels highlighting clearly delineated blood vessels from 750 – 850 μm (**Fig. 3**).

**Fig. 2.**
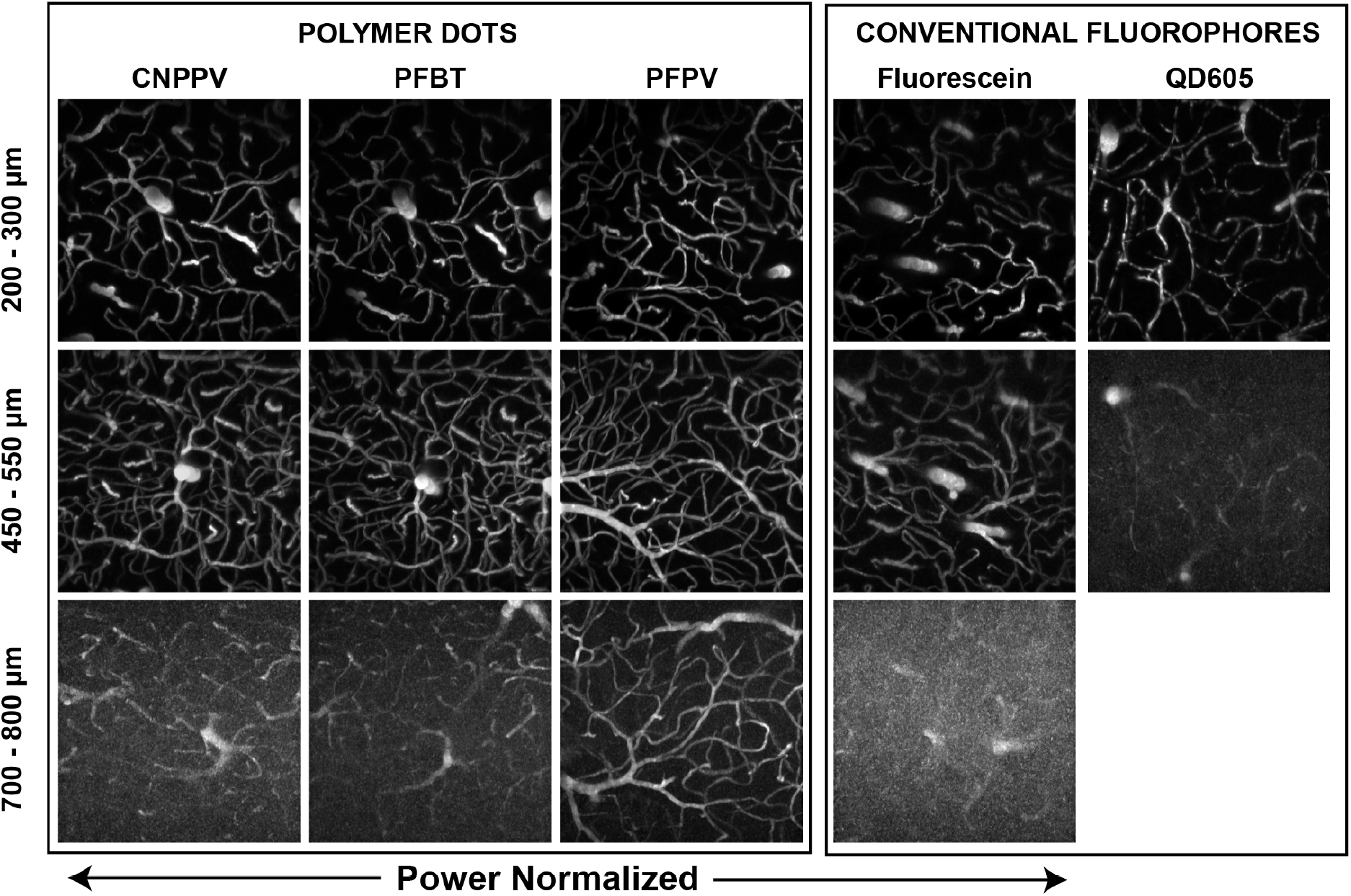
Titanium-sapphire *in vivo* vascular imaging of C57 mice labeled by a retro-orbital injection of polymer dots (*Columns 1-3*), dextran-conjugated fluorescein (*Column 4*), and semiconductor quantum dots (QD605; *Column 5*); λ_ex_ = 800 nm. Maximum intensity projections spanning 100 μm ranges were taken at various depth intervals in the cortex, with distance relative to the pial surface indicated across the left margin. Polymer dots yield appreciably higher signal-to-background ratio at all depths relative to fluorescein and QD605. Imaging of QD605-labeled vasculature beyond 550 μm was restricted due to the toxicity of the injection.

**Fig. 3.**
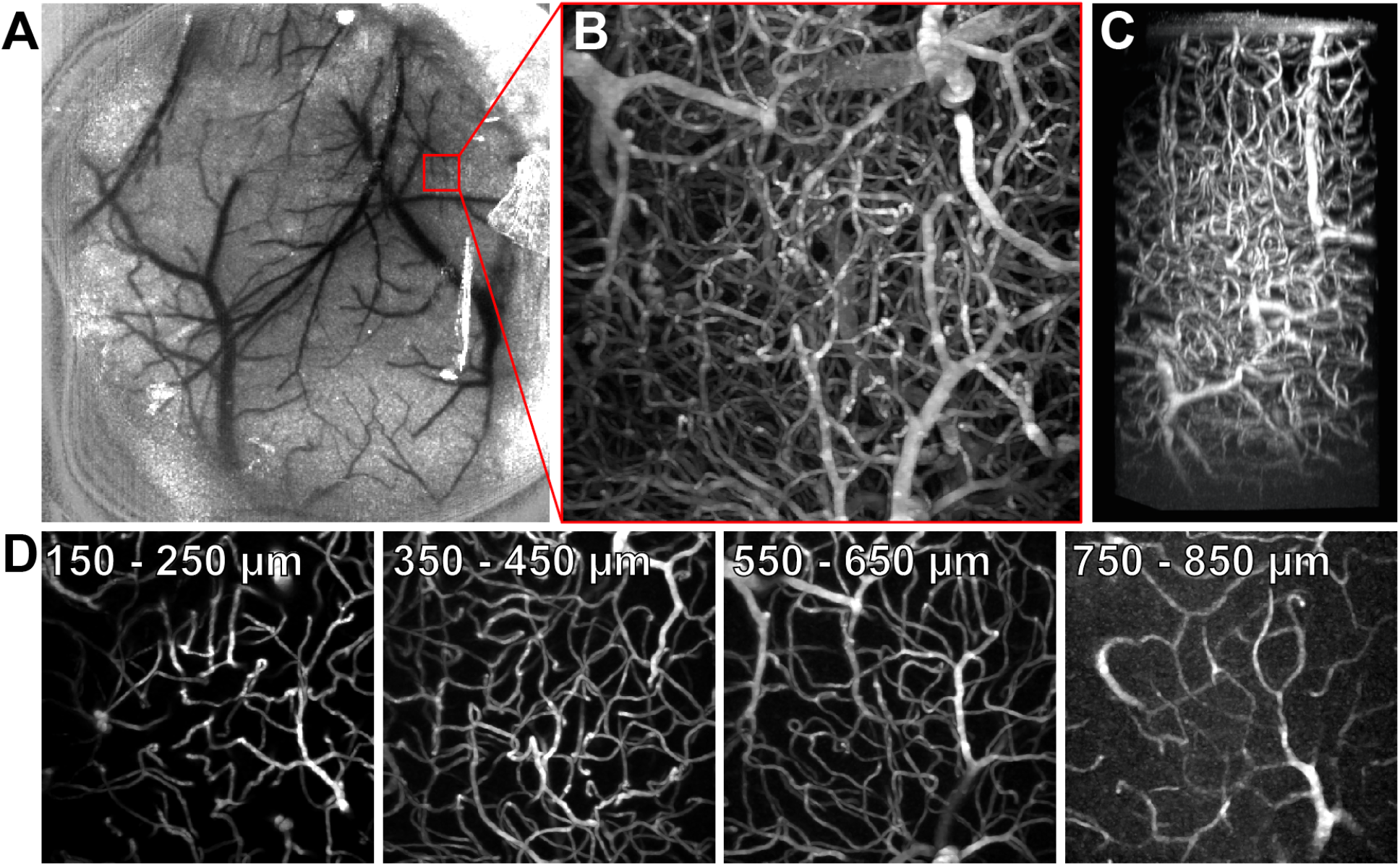
Two-photon imaging of PFPV polymer dots intravenously injected in C57 mice. *(A)* Laser speckle contrast image of surface blood vessels. *(B)* A tangential (xy) maximum intensity projection of a 365 × 365 × 850 μm image stack collected from the region of interest (ROI) delineated in red in panel A. *(C)* A max intensity projection of a 3D reconstruction from the same data. *(C)* 2D tangential (xy) projections over shorter depth ranges of the stack.

### Polymer dot excitability at longer wavelengths extends vascular imaging depth and signal-background-ratio

A distinct characteristic of polymer dots is their wide absorption range.^19^ The broad excitability of polymer dots enables their compatibility with a number of excitation sources including long wavelength, tunable laser sources between λ_ex_ = 1,100 and 1,400 nm. This allows us to employ a unique laser system optimized for deep imaging applications (**Fig. S5**) as well as excite polymer dots at longer wavelengths. The maximum imaging depth of MPM is ultimately determined by SBR, which is influenced by the scattering and absorption events that occur in biological tissue as well as the power dependence of an emitter. Excitation light that is attenuated by scattering or absorption before reaching a contrast agent fails to produce any emitted fluorescence, and the fraction of attenuated photons is heavily dependent on wavelength. The fraction of excitation light reaching the focal volume at a depth, *z*, can be approximated as exp[-(μ_a_(λ)+μ_s_(λ))*z*], where μ_a_(λ) and μ_s_(λ) are wavelength-dependent absorption and scattering coefficients.^37^ This function reveals that there is an ideal biological imaging wavelength situated at 1,300 nm^23^ where photo-attenuation is minimized in brain tissue (**Fig. S6**). In addition, 1,300 nm light can be used to achieve three-photon excitation of many fluorophores, which reduces out-of-focus excitation and thereby decreases background^9,38,39^ compared with two-photon excitation. Therefore, it is readily understood that brighter fluorophores such as polymer dots that exhibit a strong 3P power dependence in response to 1,300 nm excitation should yield a markedly improved SBR and maximize imaging depth. Unfortunately, the 3PA cross section of a polymer dot such as CNPPV is quite low at 1,300 nm. However, a suitable compromise for an optimal imaging wavelength is at 1,225 nm where attenuation length remains relatively close to its maximum (**Fig. S6**) and the slope of CNPPV’s power dependence is ~2.5 (**Fig. S2**), representing a combination of two- and three-photon excitation processes. Indeed, longer wavelength excitation of CNPPV-labeled vasculature (λ_ex_ = 1,225 nm) allows us to exploit these principles for optimized deep imaging and achieve a tissue penetration depth up to 1.3 mm beneath the pial surface (**Fig. 4A**). In contrast, two-photon imaging (λ_ex_ = 800 nm) of the same region of the same mouse only reached a maximum imaging depth of 850 μm (**Fig 4B**). A comparison of the CNPPV-labeled vascular networks imaged at the different wavelengths shows that SBR is vastly improved by 1,225 nm excitation and a higher-order power dependence (**Fig. 4C and 4D**). The effect is pronounced enough that the SBR of the 1225 nm image stack at *z* = 900 μm (SBR ~ 7.8) exceeds the SBR of the Ti:S image stack near the cortical surface (SBR ~ 7.2; *z* = 350 μm). A plot of background signal versus depth shows that the difference in SBR is directly owed to the rapid increase of background introduced by 800 nm excitation relative to the modest rise of background seen from 1,225 nm excitation (**Fig. 4D**). Quantification of SBR and background throughout the superficial dura (*z* = 0 – 200 μm) is omitted from these plots due to high background caused by second harmonic generation of collagen.

**Fig. 4.**
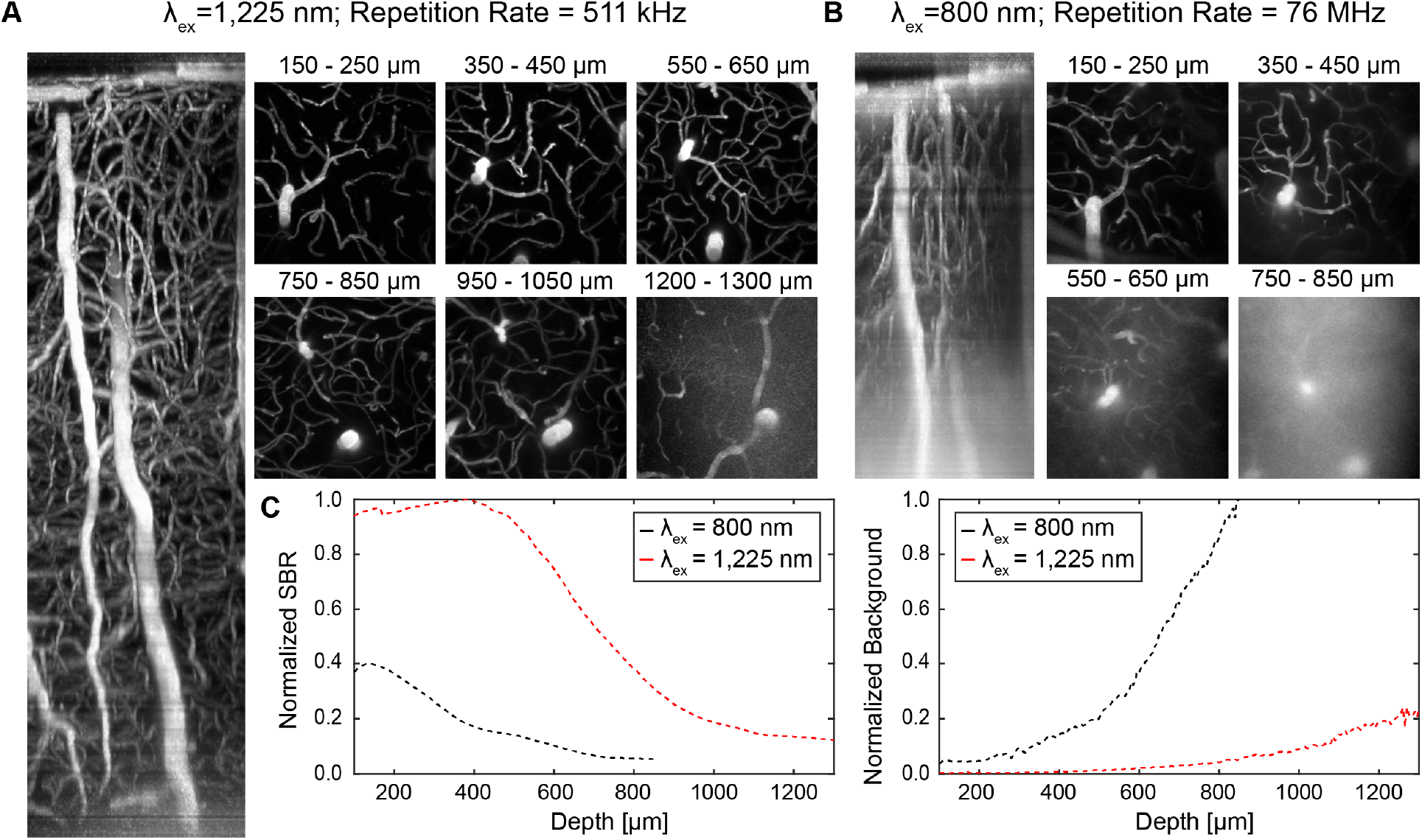
Longer wavelength excitation (λ_ex_ = 1,225 nm) versus shorter wavelength excitation (λ_ex_ = 800 nm) of CNPPV-labeled C57 vasculature. *(A)* Sagittal (*xz*) projection of a 365 × 365 × 1300 μm^3^ image stack at λ_ex_ = 1225 nm using an optical parametric amplifier source. *(B)* Sagittal (*xz*) projection of a 356 × 365 × 850 μm image stack of the same region show in *A* at λ_ex_ = 800 nm using a titanium-sapphire source. *(C)* 2D tangential (*xy*) projections over shorter depth ranges of the stacks shown in *A* and B. *(D)* Comparison plots of signal-to-background ratio (SBR) versus depth (*Left*) and background intensity versus depth (*Right*).

### High Power Fiber Laser Imaging of Polymer Dots Significantly Improves Signal-To-Background Ratio Beyond ~600 μm Depth

Polymer dots’ broad light absorption further permits excitation by a high average power ytterbium-fiber laser (λ_ex_ = 1,060 nm). Fiber lasers are a lower-cost alternative to OPA imaging, and offer excellent pulse characteristics as well as ease-of-operability to the user.^40^ Our lab recently has recently demonstrated the use of such a system for deep *in vivo* imaging.^6^ Although the fiber laser’s fixed output wavelength (λ_ex_ = 1,060 nm) does not coincide with the ideal biological imaging wavelength (λ_ex_ = 1,300 nm), a comparison of *in vivo* image stacks of PFPV-labeled C57 vasculature collected at λex = 1,060 vs 800 nm clearly illustrates the advantages of the polymer dots’ excitability by a fiber laser (**Fig. 5**). Although resolvable imaging depth is not significantly extended by the yb-fiber (*z* = 900 μm) relative to the Ti:S penetration limit (*z* = 850 μm), the image quality (contrast and SBR) of the yb-fiber images is substantially improved under 1,060 nm excitation, an effect which is most prominent beyond ~600 μm (**Fig. 6A**). This corresponds to the *z*-position at which maximum Ti:S output power was reached whereas the upper limit of fiber laser output power was not met before all signal was lost. In addition to the higher average power of the yb-fiber relative to the Ti:S source, two primary advantages can be attributed for the dramatic improvement in SBR. First, the number of photons lost to scattering and absorption events is reduced by longer wavelength excitation (λ_ex_ = 1,060 nm vs. 800 nm) (**Fig. S6A**). At an 850 μm depth, the remaining photon fraction at λex = 1,060 nm is approximately five times greater than at λ_ex_ = 800 nm. Second, PFPV exhibits a partial three-photon power dependence at 1,060 nm (n ~ 2.38 ± 0.08) versus a two-photon excitation signature at 800 nm (n ~ 1.93 ± 0.14) (**Fig. S2**). This partial three-photon dependence at λ_ex_ = 1,060 nm further improves SBR through suppression of background fluorescence due to a higher-order nonlinear dependence on excitation intensity. Thus, longer wavelength excitation of PFPV is expected to improve both the signal and background of images at all depths.

**Fig. 5.**
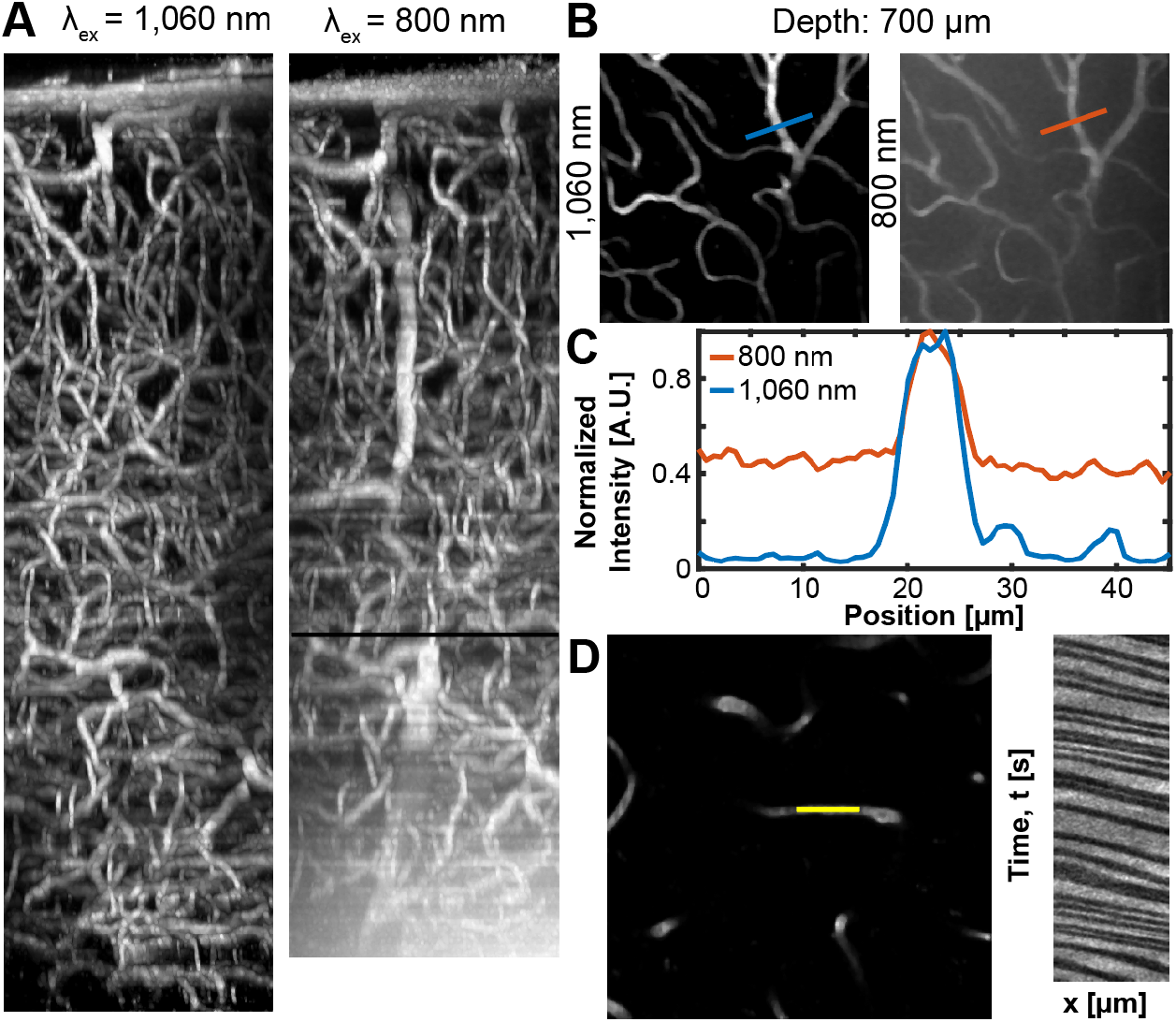
High power fiber laser imaging of PFPV-labeled vasculature improves the SBR of images. (A) Image stacks of the same cortical region collected using a 1,060 nm ytterbium-fiber laser (*left*, 256 x 256 x 900 μm^3^) or a 800 nm Ti:S excitation source (*right*, 256 x 256 x 850 μm^3^). The depth at which maximum power output from the Ti:S laser was reached is marked by a black line (*z* = 600 μm). (B) 50 μm thick maximum intensity projections of the images stacks shown in *A* centered at 700 μm. The blue and red lines denote the positions of analyzed 45 μm long line profiles. *(C)* A plot of normalized signal intensity relative to position. (*D*) A vessel line scan collected at a depth of 750 μm. The analyzed blood flow velocity is 1.29 ± 0.20 mm/sec.

This expectation is empirically demonstrated by line profiles drawn across identical in-plane blood vessels located 700 μm beneath the surface imaged separately at λex = 1,060 and 800 nm (**Fig. 5B-C**). The data reveals that the SBR of the λ_ex_ = 1,060 nm image is ~3.5 times greater than the λ_ex_ = 800 nm SBR. Dramatic SBR improvement at z-positions beyond the depth at which Ti:S excitation becomes power-limited demonstrates that the higher output power of the yb-fiber laser coupled with longer wavelength excitation is a critical advantage for deep *in vivo* imaging. We emphasize that similar improvements made by longer wavelength excitation cannot be achieved with conventional fluorophores such as fluorescein (*σ*_2_ = 0.31 at λ_ex_ = 1,050 nm) or Texas Red, which exhibit low action cross sections beyond the Ti:S range (>1,000 nm) and are unlikely to undergo a three-photon transition at 1,060 nm.^5,27^ A major practical benefit of improved SBR is the ability to collect high-quality vessel line scans at greater depths to quantify blood velocity. Vessel line scanning quantifies flow velocity along the central axis of blood vessels at high frequency, relying on the contrast between injected plasma fluorophores and red blood cells (RBCs) which remain dark.^41^ Therefore, high SBR is essential to accurately differentiate RBCs and fluorescent plasma and obtain precise flow velocity measurements. Here, we show that vessel line scans of PFPV-labeled mice form an image containing distinct streaks that represent RBCs traveling at a mean velocity of 1.28 ± 0.20 mm/sec at a depth of 750 μm beneath the pial surface (**Fig. 5D**).

## Discussion

To evaluate polymer dots’ multiphoton properties, we prepared multiple polymer variants via nanoprecipitation, a simple and rapid procedure. The batch size could be easily varied to prepare ample amounts of the nanopolymers for vascular imaging of tens of mice per preparation. The polymer dots had the additional advantage of a long shelf life (upwards of 5 months) and easy storage, characteristics that allow them to be easily disseminated for broad use.^42^ Through our imaging experiments, we demonstrate that polymer dots offer a wealth of optical properties that make them very well suited for deep, *in vivo* multiphoton fluorescence microscopy. However, the lack of knowledge surrounding their nonlinear excitation properties has prevented their widespread adoption as popular contrast agents in multiphoton fluorescence imaging. To remedy this, we have characterized the two-versus three-photon power dependence of three polymer dot variants (CNPPV, PFBT, and PFPV) representing different polymer classes including poly(phenylene vinylene) and fluorene-based copolymers (**Fig 1**). We learn that all three polymer dot species demonstrate a 2P power dependence across the conventional Ti:S tuning range (λ_ex_ = 780 – 850 nm). Meanwhile, researchers who rely on the photophysical advantages that accompany 3PM, such as reduced out-of-focus fluorescence and diminished scattering, in order to improve the signal-to-background ratio in their imaging experiments can do so via excitation of CNPPV at 1300 nm, PFBT at 1350 nm, and PFPV at 1325 nm. This is a stark contrast to conventional organic dyes such as Texas Red and ICG, which remain rooted in the two-photon regime at comparable wavelengths (λ_ex_ = 1,350 nm and 1,280 nm, respectively).^5^ Furthermore, it is interesting to note that CNPPV undergoes a three-photon transition at a higher energy wavelength than either PFBT or PFPV, despite being the most red-shifted of the three. This can potentially be explained by the unique chemical structure of poly(phenylene vinylene) polymers relative to fluorene-based co-polymers. The distinct molecular structures of the conjugated polymers affect the final energy levels of the polymer dot species in their excited states. Specifically, CNPPV has a donor-π-acceptor arrangement whereas PFBT and PFPV have donor-acceptor and donor-π-donor configurations, respectively, which leads to differences in the energy levels of their three-photon transitions.

Furthermore, we justify the use of polymer dots over conventional fluorophores for multiphoton imaging experiments in the 800 – 900 nm excitation range due to their extraordinary brightness relative to quantum dots, organic dyes, and fluorescent proteins (**Fig. S3**). PFPV (~2.02×10^5^ GM) is brighter than MEH-PPV (~5.50×10^4^ GM) and their 2PA cross sections are one to two orders of magnitude larger than an expected range for quantum dots (~2×10^3^ to ~4.7×10^4^ GM). When compared to fluorescent dyes and proteins, polymer dot cross sections are three to five orders of magnitude larger. Most notably, this comparison includes organic molecules specifically engineered for enhanced two-photon absorption.^28,43^ In particular, many fluorescent dyes can be problematic within *in vivo* biological settings due to a reduced quantum yield and poor photostability in aqueous environments.^13–15^ In contrast, polymer dots composed of hydrophobic conjugated polymers are strongly resistant to this effect.^19^ Of course, highly emissive quantum dots such as QD605^28^ and QD535^27^ do come close to matching a dimmer polymer dot such as MEH-PPV in brightness. Therefore, one might be tempted to employ quantum dots as their preferred class of contrast agents due to their well-regarded photostability and spectrally broad light absorption, an essential feature for multicolor imaging experiments. However, polymer dots share these same properties^20^ as well as the critical advantage of exhibiting no evidence of cytotoxicity. There are conflicting reports regarding quantum dot biocompatibility which can most likely be attributed to physiochemical and environmental factors;^18^ yet in our hands, intravenous injection of QD605 was acutely toxic and fatal to mice whereas identical chronic studies with polymer dots did not produce any observed health concerns. Nevertheless, a biological mechanism of polymer dot blood clearance has yet to be thoroughly assessed and future research efforts should be directed to determine whether bioaccumulation in the spleen, liver, kidney or other major organs is a valid concern.

Next, we were able to show that polymer dots’ large action cross sections and excitability in the NIR regime improves SBR at considerable depths in mouse cortex at an 800 nm excitation wavelength relative to identical experiments performed with an exceptionally bright quantum dot species, QD605, and a commonly used organic dye, dextran-conjugated fluorescein (**Fig. 2**). As noted earlier, QD605 is cytotoxic when administered intravenously, particularly at the high concentrations necessitated by deep imaging experiments. *Ex vivo* animal studies may not be hindered by such a consequence; however, in the case of vascular imaging, continuous circulation and blood flow is essential to avoid distorted vessel appearance and photobleaching of the now stagnant probes (**Fig. 2**, *right*). The overall imaging depth of the fluorescein-labeled vasculature matched that of the polymer dot-labeled mice; however, the SBR of the fluorescein images were less satisfactory resulting in reduced vascular clarity (~1.8-fold at *z* = 550 μm; **Fig. S4**). The implications of poor SBR include obfuscated analysis and degradation of automated image segmentation schemes.^44^

Another substantial limitation of fluorescein and other conventional organic dyes is their narrow absorption range. At λ_ex_ = 800 nm, fluorescein’s 2PA action cross section is ~36 GM and at λ_ex_ = 1,050 nm its cross section is a barely detectable 0.31 GM.^27^ In contrast, polymer dots exhibit broad multiphoton absorption, meaning that they can be efficiently excited with a variety of laser sources all the way out to 1,400 nm. We were able to demonstrate that the use of longer excitation wavelengths and higher-order nonlinear excitation allows us to improve SBR and overcome the tissue scattering limits of imaging depth observed with conventional 2PM. Specifically, excitation at λ_ex_ = 1,225 nm attains a 1,300 μm imaging depth whereas λ_ex_ = 800 nm results in an 850 μm cortical depth (**Fig. 4**). Moreover, the SBR of the images collected at the longer wavelength excitation greatly exceed those recorded at an 800 nm excitation at all depths, producing a much higher quality 3D volume. Through modeling (**Fig. S5**) and our power dependence characterization (Fig. S2) we are able to attribute this gain in penetration depth and SBR to the reduced scattering and photo-attenuation of longer excitation wavelengths, and the ancillary reduction of background signal due to CNPPV’s partial 3P power dependence at 1,225 nm.^45^ In regard to the neuroscience community, the extended imaging depth with longer wavelength excitation enables researchers to investigate neural layers of anatomy beyond the cortex and corpus callosum, most notably the hippocampus. This holds exceptional significance due to the role of hippocampal hemodynamics in neurogenesis and severe cognitive pathologies such as Alzheimer’s disease.^46,47^ In addition, the biological imaging community can take advantage of polymer dots’ overlapping broad absorption spectra, which permits several polymer dot species emitting at discrete bands to be excited simultaneously at a single wavelength. This characteristic coupled with the fact that polymer dots can be readily functionalized and used to tag unique cellular structures^34^ allows researchers to design simple and effective multicolor imaging experiments. An important caveat to consider, however, is that polymer dots were delivered intravenously in our studies, and labeling neural structures located in high-density extravascular brain tissue could pose a challenge due to the relatively large diameters of polymer dots (~20-30 nm). Recent efforts have produced polymer dot nanoparticles with sub-5 nm diameters, yet the yield from these preparations is still quite low.^48^

We also take advantage of polymer dots’ broad light absorption to image PFPV-labeled mice and resolve vasculature using a custom, home-built ytterbium-fiber laser.^6^ Again, we observe that reduced scattering of longer wavelength excitation light and a partial 3P power dependence improves imaging depth slightly (*z*_800 nm_ = 850 μm; *z*_1060 nm_ = 900 μm) (**Fig. 5**). We note that the SBR of the 1060 nm excitation images is improved most substantially beyond ~600 μm, an imaging depth which corresponds to the *z*-position at which the Ti:S output power was saturated.

Although we were able to significantly improve SBR and extend imaging depth using longer wavelength excitation (λ_ex_ = 1,060 nm and 1,225 nm) we were unable to achieve similar results when imaging at an ideal biological imaging wavelength identified through modeling (λ_ex_ = 1,300 nm) due to polymer dots’ lower brightness at this wavelength. Advances in polymer engineering, structure based-design, and mutagenesis can vastly improve polymer dot performance and enhance multiphoton absorption at 1,300 nm for optimized deep imaging. Future redesign and optimization of polymer dots’ molecular structure for improved three-photon imaging will involve an exploration of the effects of donor-π-donor, donor-π-acceptor-π-donor, and acceptor-π-donor-π-acceptor conformations on absorption cross sections and quantum yield. Overall, polymer dots present an exciting new approach to multiphoton *in vivo* imaging due to their enhanced brightness, broad excitability, and nontoxic features. With brighter, biocompatible probes, researchers will be able to resolve vascular architecture in living organisms with improved clarity and depth, enabling critical insights into fundamental biological problems.

## Material and Methods

### Power Dependence Measurements

Adjacent sample wells in a multi-channel slide (μ-Slide VI0.4, Ibidi) were filled with 40 μl of 100 nM CNPPV, PFBT, or PFPV alongside a blank solution (0.9% blood bank saline, Fisher). The sample was then exposed to laser light excitation and the emitted photon counts were monitored by a digital photon counting board (DPC-230, Becker & Hickl GmbH). Three consecutive measurements of photon counts from the sample and blank were recorded using a five second acquisition duration in absolute timing mode with a threshold of −120. Signal was maximized immediately before each recording by adjusting the *z*-position of the sample surface relative to the objective via a motorized lab jack (L490MZ, Thorlabs). Measurements were repeated at increasing power intensities and the logarithmic values of the background-subtracted signal were plotted as a function of the logarithmic excitation value. A linear model was then fit to the data (MATLAB, National Instruments) and the slope was extracted to determine the power dependence. A fresh sample was used for each distinct wavelength to avoid the potential introduction of any artifacts related to photostability. Excitation wavelengths of the Ti:S source (Mira 900) and the yterrbium-fiber laser (home-built) were measured with a UV-Vis-NIR spectrometer (USB2000, Ocean Optics) whereas the longer optical parametric amplifier wavelengths were measured with an NIR spectrometer (AvaSpec-NIR256-1.7, Avantes).

### Two-photon microscopy

In the two-photon laser schematic, a 10W Verdi is used to pump a mode-locked titanium:sapphire (Ti:S) oscillator (Mira 900, Coherent). The beam is steered to a pair of galvanometer scanners (6125HB, Cambridge Technology) driven by servo driver amplifier boards (671215H-1HP, Cambridge Technology). A Keplerian telescope beam expander which consists of a B-coated scan lens (f = 80.0 mm, AC254-080-B, Thorlabs) and tube lens (f = 200.0 mm, LA1979-B-N-BK7, Thorlabs) is used to properly fill the back aperture of the microscope objective (XLUMPLFLN20XW 0.95 NA or XLPLN25XSVMP2 25X 1.0 NA, Olympus). Excitation and emission paths are separated with a 775 nm cutoff dichroic mirror (FF775-Di01-52x58, Semrock). Fluorescence is epi-collected, transmitted through either a 510/84 bandpass filter (FF01-510/84-25, Semrock) or a 609/181 bandpass filter (FF01-609/181-25, Semrock), and detected by a photomultiplier tube (H10770PB-40, Hamamatsu Photonics). The current signal is then sent to a current pre-amplifier (Model SR570, Stanford Research Systems) and digitized by a PCI-based data acquisition (DAQ) board (PCI-6353, National Instruments). Image acquisition was controlled using custom software (LabVIEW, National Instruments) and image frames were collected at a 512 x 512 pixel size. Image stacks were collected at a *z*-resolution of 5 μm and three frames were averaged from 0 – 200 μm cortical depths, five frames from 200 – 500 μm, eight frames from 500 – 700 μm, and twelve frames beyond 700 μm. All mice specimens imaged by Ti:S were excited at λ_ex_ = 800 nm.

### Three-photon microscopy

The three-photon laser schematic is shown in **Supplementary Figure 5**. A 5W laser is used to pump a λ_ex_ = 800 nm mode-locked Ti:S oscillator (Mira 900, Coherent) which is stretched by an external stretcher/compressor to seed a regenerative amplifier (RegA 9000, Coherent). The amplified pulse is compressed and the spectrum of this 800 nm pulse can then be shifted and tuned by an optical parametric amplifier (OPA 9800, Coherent). Three-photon *in vivo* imaging was performed at λ = 1225 nm using a 25x multiphoton objective (XLPLN25XSVMP2 25x 1.0 NA, Olympus). Again, image stacks were collected at a *z*-increment of 5 μm and all frames were 512 x 512 pixels. Three frame running averages were recorded for the first 500 μm and an additional two frames were averaged each 200 μm interval beyond that.

### Fiber Laser Microscopy

A commercial fiber oscillator (Origami-10, OneFive GmbH) was used to seed a custom-built ytterbium fiber amplifier. The amplifier consisted of a 6-meter long segment of double-clad ytterbium-doped polarization-maintaining large-mode-area optical fiber with a core diameter of 25 μm and cladding diameter of 250 μm (YB1200-25/250DC-PM, Thorlabs), coiled to a radius of less than 7 cm to suppress higher-order modes and achieve single-mode output. The fiber was pumped by a fiber-coupled laser diode emitting up to 30 W of 915-nm light (K915FA3RN-30.00W, BWT Beijing. With a pump power of 23W, the amplifier produced 6 W of output at 1060 nm with a repetition rate of 80 MHz, and a grating pair was used to compress the pulse width to ~120 fs in the sample plane. The laser power was adjusted using a Glan-type calcite polarizer/half wave-plate combination.

### Vessel Line Scanning

Line scans were acquired along vessel axes by collecting the fluorescence intensity while the galvo mirrors rapidly scanned a linear, user-defined path along a capillary. Each line consisted of 200 pixels imaged at a rate of 450 kHz, enabling a line rate of 940 Hz. To build a trajectory, the fluorescence intensity is plotted as a function of time (line number) and space (xy-coordinate, or pixel in one line), as shown in Figure 5(D). A sufficient number of lines were scanned along a trajectory to quantify the blood flow from analyzing the dark streaks. A Radon transform was used to determine the inclination angle of the dark streaks which, when combined with the known temporal and length scales, allowed for calculation of the RBC velocity.^41,49,50^

### Image Analysis

To quantify signal to background ratio, a custom script (MATLAB, National Instruments) was used to read in an image sequence, reshape the image intensity matrices into one-dimensional arrays and sort the pixel values in ascending order. The top 10% of cells were averaged to obtain a signal value and the bottom 70% of cells were averaged to obtain a background value, from which a signal-to-background ratio could easily be calculated. To obtain intensity profiles, images were loaded into image analysis software (ImageJ, NIH) and the line selection tool was used to demarcate a region of interest. A plot profile was then extracted to display a two-dimensional graph of pixel intensities versus position.

## Funding Acknowledgments

This study has been supported by the National Institutes of Health [NS082518, NS078791, EB011556 CA193038], the American Heart Association [14EIA18970041], and the Texas 4000 Foundation and Cancer Prevention Research Institute of Texas [RR160005]. Daniel T. Chiu gratefully acknowledges support of this work by the NIH [MH113333].

